# The double life of trichomes: understanding their dual role in herbivory and herbicide resistance

**DOI:** 10.1101/2023.04.28.538721

**Authors:** Nia M Johnson, Regina S Baucom

## Abstract

External features of organisms often serve as the first line of defense in their immediate environments. Trichomes are hair-like appendages on plant surfaces that can defend against damaging agents such as pathogens, herbivores, and UV radiation. It is currently unknown if the variation observed in trichome traits represents dual or conflicting roles against different types of stressors. Here, we assess whether trichomes serve as an herbicide resistance trait and how it coincides with the conventionally studied defensive strategy of herbivory resistance. In a series of experiments, we exposed the annual invasive velvetleaf (*Abutilon theophrasti*) to glyphosate (active ingredient in the herbicide ‘Roundup’) to investigate whether trichome traits (shape and density) are linked to herbicide resistance and to test whether herbicide alters selection on plant trichomes. We found that an increased proportion of branched trichomes positively impacted herbicide resistance as well as chewing herbivory resistance. We also found evidence that glyphosate imposes positive selection on branched trichomes in velvetleaf. Overall, our results indicate that branched trichomes can contribute to both herbicide and herbivory resistance, serving as a dual structural form of resistance reducing plant injury. If our results are to be applied more broadly, it would suggest that herbicide exposure can alter the composition of plant trichomes, potentially impacting trichome-mediated defenses against various external stressors.

## Introduction

Organisms cope with many types of external stressors, which can lead to wide variation of defensive traits. Often, this trait variation is a result of complementary or conflicting strategies for dealing these stressors (Simms 1992, Koricheva 2001, Fornoni et al 2004) and the relationships between defense traits and fitness in any given environment. Of the various types of defense strategies in plants (chemical toxicity, low digestibility, leaf toughness, etc.) one of the most important may be the multifaceted traits that make up the landscape of the physical boundary, since, once encountered, this boundary acts as the first line of defense and plays a critical role in how plants interact with their environment.

A key form of defense found across a range of plants on the outermost boundary are hair-like appendages referred to as trichomes (derived from the Greek word for hair: trichos). These unicellular or multicellular structures can create complex networks that respond to external cues (Werker 2000). While traditionally viewed as a defensive strategy against herbivory (Levin 1973, Stipanovic 1983, Southwood 1986), trichomes also play a protective role against other biotic stressors such as pathogens (Shepard et al 2005, Gao et al 2017) and abiotic stressors such as UV-B radiation, ground-level ozone, and drought (Yan et al 2012, Li et al 2018, Bickford 2016). Leaf trichomes have also been shown to play a role in the detoxification of heavy metals by way of compartmentalization (Küpper et al. 2000, Marmiroli et al 2004, McNear 2013) and secretion (Psaras et al 2000).

The many potential functions of trichomes likely results from different selection pressures acting simultaneously on trichome phenotypes, including their morphology and density. For example, trichomes may serve as an herbivory resistance trait, but the evolution of a particular trichome phenotype that optimizes herbivory resistance may come at the cost of increased susceptibility to other agents of damage like UV, drought, or herbicide. In this case, trichome phenotypes would play a ***conflicting*** role in plant defense, as selection for one function (*i*.*e*., to reduce the fitness effects of herbivory) may contradict a different function leading to increased susceptibility to another type of damage (*i*.*e*., increased susceptibility to herbicide).

Alternatively, selection for increased resistance to one agent of damage may likewise lead to increased resistance to a different agent of damage, thus serving a ***dual*** defensive role (Agrawal and Fishbein 2006).

Here, we evaluate the dual or conflicting role that trichomes may serve when defending a plant to multiple stressors. The agents of damage we consider are herbivory and herbicide, both of which natural populations of plants regularly experience in agricultural ecosystems. We do so using the weed *Abutilon theophrasti*, commonly referred to as velvetleaf because of the dense layer of trichomes that blanket the surface of its leaves. This species is known for being covered in distinct polymorphs of trichomes (branched, single, peltate, and capitate; Sterling 1978), which may each play different functional roles in plant defense (Levin 1973). Furthermore, the species has evolved reduced susceptibility to multiple herbicide classes, including triazine and glyphosate (Anderson and Gronwald 1991, Hartzler and Battles 2001), but the mechanisms underlying this reduced susceptibility are presently unknown. While leaf surface morphology can influence herbicide distribution and absorption (Hess and Falk 1990, Hess 2018), the most well-studied herbicide resistance mechanisms are biochemical rather than structural (reviewed in Gaines et al 2020). Thus, despite their implication as a possible herbicide resistance mechanism (Devine et al 1992, Baucom 2019), it remains unknown if trichomes can serve as a form of structural defense against herbicide. Furthermore, just as trichome traits are subject to selection from herbivory (Valverde et al 2001) and UV or drought (Steets et al 2010, Tang et al 2020), herbicide may act as an agent of selection on trichomes. To our knowledge, however, this hypothesis has never been formally tested by examining selection on trichome traits in the presence and absence of herbicide.

To test whether trichomes are undergoing conflicting or parallel selection from different stressors, we ask the following questions: 1) Do trichomes serve as a defense trait to herbivory and/or herbicide in *A. theophrasti*? A positive correlation between trichome traits and either herbicide or herbivory resistance would provide evidence that trichomes serve as a defense trait to either respective agent of damage. We next ask, 2) Do trichomes perform a dual or conflicting defense role given exposure to herbicide and herbivory? A positive correlation between trichomes and both herbivory and herbicide resistance would provide evidence that trichomes serve a dual defensive role, whereas a positive correlation with one form of resistance but a negative correlation with the other would suggest a conflicting role. Finally, we ask 3) Does herbicide act as an agent of selection on trichome traits? Evidence of positive selection on a particular trichome phenotype would indicate that herbicide is selecting for that phenotype, whereas negative selection would indicate that herbicide is selecting against that phenotype.

While manipulative experiments in other plant systems show convincing evidence that plant herbivores act as selective agents on trichomes (Mauricio and Rausher 1997), to our knowledge no such relationship has been established for trichomes and herbicide.

## Methods

### Study Organism

*Abutilon theophrasti* Medicus, (Malvaceae) is an invasive annual native to Asia and is frequently found in and around corn (*Zea mays*) and soybean (*Glycine max*) fields. In contrast to its historic past as a fiber crop for early North American colonists, *A. theophrasti* is presently a major weed species in North America as well as other temperate regions between 32º and 45º N latitude.

Studies in South Dakota (Scholes et al 1995) and Pennsylvania (Werner et al 2004) suggest that velvetleaf can account for maximum annual losses of 33% and 37% on corn and grain, respectively. Annual herbicide cost of this species has been estimated at $55.00/ha in corn fields alone (Werner et al 2004), with an estimated 32.75 million ha of corn fields being treated with herbicide within the US (ISAAA, 2017). The success of this competitive ruderal is largely due to reproductive plasticity including self-pollination capabilities (Warwick and Black 1985), prolific seed production (can produce >8,000 per plant), seed dormancy for 50+ years (Spencer 1984), and reduced susceptibility to herbicides (Sattin et al 1992).

*Abultion theophrasti* gets its common name, velvetleaf, from the velvety feeling produced by a soft, dense layer of trichomes on the plant surface. Velvetleaf produces four trichome types. Two are multicellular, glandular forms that synthesize and secrete metabolites. Two are unicellular and non-glandular trichomes. The glandular types include peltate trichomes, which are globular structures composed of 4-5 cells, and capitate trichomes, which are stalked structures composed of 12-15 cells. There are two non-glandular trichomes – single and branched, which are structures that grow perpendicular to the plant surface, or single cell structures consisting of 4-8 arms, respectively (Figure 1). As a species that possesses the major trichome classifications observed in nature, velvetleaf is a powerful system to study the role of trichome-mediated defense.

**Figure 1.**
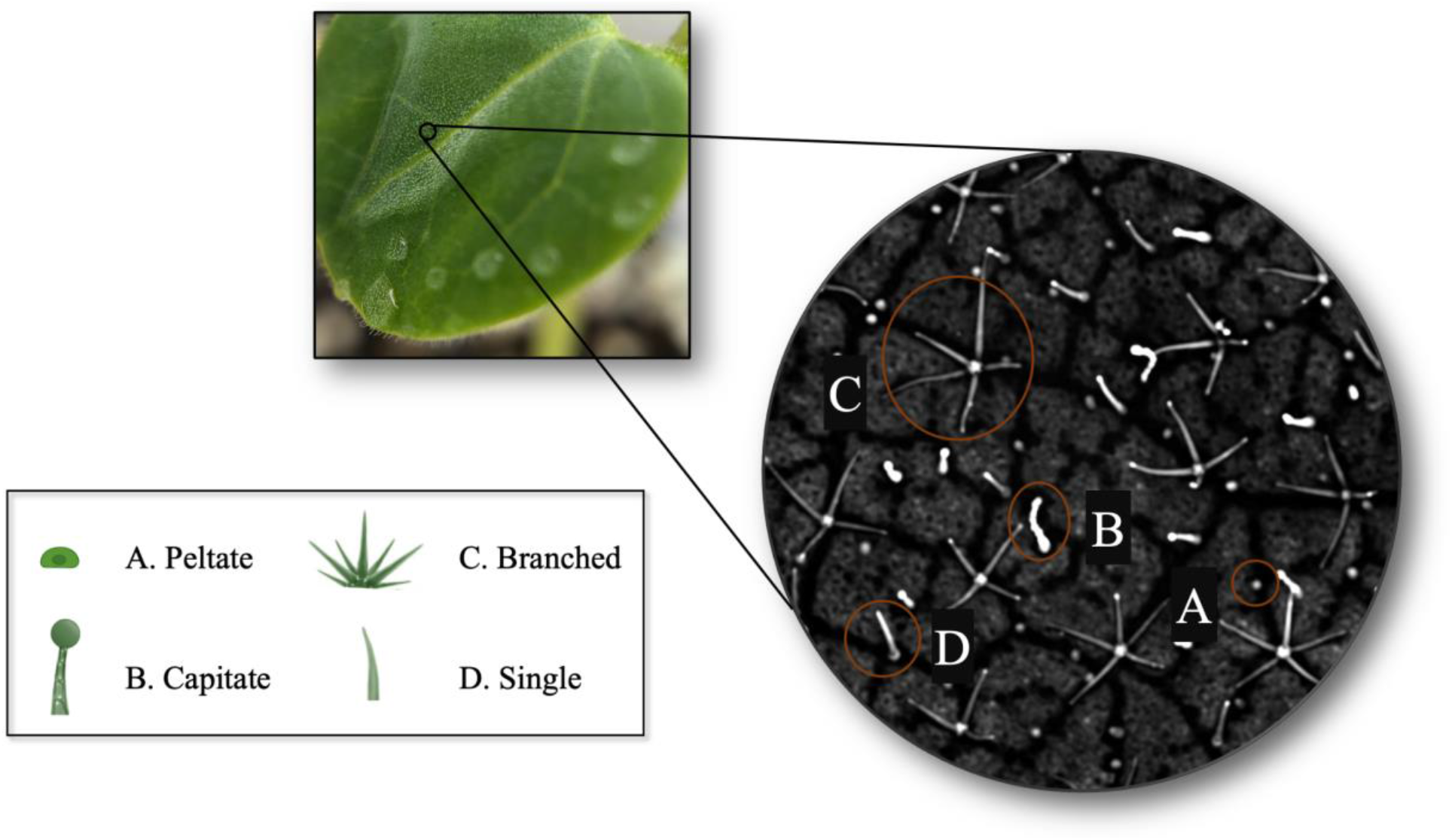
Characterization of the trichomes present on adaxial surfaces of *Abutilon theophrasti* leaves using confocal imaging (x10). A) Peltate trichomes are multicellular glandular structures made up of 4-5 cells. B) Capitate trichomes are multicellular glandular structures made up of 12-15 cells. C) Branched trichomes are single cellular structures. D) Single trichomes are unicellular unbranched structures. Density is calculated as the total number of all trichomes captured in the image.

We consider trichome phenotypes in two main ways – first and following typical evolutionary ecology studies (Kauffman and Kennedy 1989, Soetens et al 1991, Gianfagna et al 1992, Strauss et al 2003, Castillo et al 2013, Zhou et al 2018), we measured the total density of trichomes present on the adaxial surface of the leaf. Second, we examined the proportion of trichome types (branched, single, peltate, capitate) present on the adaxial surface to account for potential investment in different trichome types across individuals. We characterized the level of herbicide (glyphosate) resistance across *A. theophrasti* populations in the growth room and used this information to determine the appropriate herbicide dose to apply in a field experiment. In the field, we assessed the potential for correlations between trichome phenotypes (total density, proportion of each trichome type) and the level of herbicide and herbivory resistance and examined the pattern of selection on trichome phenotypes in the presence and absence of herbicide.

### Growth Room Experiment

To evaluate the level of resistance among natural populations, we collected seeds from eight velvetleaf populations in the fall of 2017 from various soybean fields around Dexter, MI (SFigure 1), and conducted a growth room experiment in the spring of 2019 using these field collections. The growth room design consisted of 6 treatments, including five levels of glyphosate (550g/ha ai, 770g/ha ai, 1125g/ha ai, 2000 g/ha ai, and 3000 g/ha ai (N- (phosphonomethyl) glycine3) Roundup® Herbicide (RU)^1^ (Monsanto Company, St. Louis, MO) and water as the control. We scarified and planted 3,120 total seeds in a randomized design of 164 ml cone-tainers (1.5in diameter, Stuewe and Sons inc., Tangent Oregon) in the growth room (130 maternal lines x 6 treatments x 4 replications) with each maternal line randomized within each treatment. We rotated plants within flats weekly. Plants were kept on a 12/12 hr light-dark regime at a temperature of 25°C. Four weeks after seed germination, when the average plant height was 11 cm tall, we sprayed glyphosate on individuals using a handheld CO_2_ sprayer, ensuring adequate coverage in each respective treatment. We harvested each plant six weeks after herbicide application to evaluate survival and the median lethal dosage (LD50), defined as the dosage of glyphosate required to kill half of the individuals for each population. To estimate herbicide resistance, we dried samples at 70°C for 3 days to get a measurement for dried biomass, a common metric for measuring herbicide resistance (Jacobs et al 1988, Vila-Aiub et al 2009, Kumar and Jha 2015).

### Field Experiment

In the summer of 2019, we conducted a field experiment to determine if there was a relationship between trichome traits and herbicide or herbivory resistance and to examine whether herbicide exposure alters the pattern of selection on trichomes. We generated seeds for this experiment by self-pollinating 40 maternal lines from the populations used as controls in the growth chamber experiment. We scarified and planted these seeds in a randomized block design with each treatment randomized within each block, and three replicate plants per maternal line present within each treatment/block combination (40 maternal lines x 2 treatments x 3 blocks x 3 replicates = 720 individuals total). Five weeks after seed germination, when the average plant height was 11 cm, we applied glyphosate at 550 g/ha ai to the glyphosate treatment, as the growth room experiment revealed the most genetic variation at this application dose. This herbicide dosage was informed by the growth room study in which the 550 g/h ai treatment exhibited the most genetic variation between populations (see SFigure 2). We collected leaves from a subset of plants for the trichome analysis (see below) and collected seeds from all plants at the end of the field season to estimate relative fitness.

### Trichome Phenotyping

We sampled the adaxial surface on one leaf from 2-3 randomly chosen plants from each maternal line and each treatment environment (herbicide/no herbicide) one week after flowering began (n = 130). We immediately brought the leaves back to the laboratory where they were preserved at 3.3°C until processing. Using a confocal laser scanning microscope (TCS SP5; Leica Microsystems CMS GmbH, D-68165 Mannheim, Germany), we captured images of leaf trichomes and quantified traits from a 1cm diameter subsection collected from the lower right quadrant of each leaf. We used a Plan Apochromat 20x/0.75 objective lens with an excitation wavelength of 488 nm and 512 × 512-pixel resolution. The confocal laser scanning microscope produced 2D image slides at 1μm increments on the z-axis plane of focus. We reconstructed these 2D sections into 3D illustrations and analyzed these images using imaging software, Fiji ImageJ version 3.0 (Schindelin et al. 2012). We applied a macro plug-in we created to recognize trichome structures and manually took counts of each type of trichome structure. Subsequently, we calculated the proportion of each trichome type by dividing the absolute number of the structure by trichome density (*i*.*e*., total number of trichomes per 1 cm diameter segment). Because raw values of plant traits are more likely to be affected by a variety of factors such as size, age, and environmental conditions not captured in our study, we elected to present the proportions of trichome types rather than the raw counts as they provide a more reliable indicator of adaptive strategy. For example, the ratio of leaf area to plant biomass is a better indicator of plant photosynthetic capacity than the absolute leaf area or plant biomass alone. Here, we report total trichome density and the proportions of trichome polymorphs to understand the functional role of trichome traits as it captures the overall landscape pattern on the leaf surface.

### Herbicide Damage

We measured herbicide damage in the field as the number of yellowing leaves two weeks following herbicide application and converted this to the proportion of the plant damaged by dividing the number of damaged leaves by the total number of leaves. We choose this metric for herbicide resistance for the field experiment because velvetleaf begins to release its leaves during seed maturation, thus it was not feasible to gather both biomass and fitness data.

### Herbivory Damage

To estimate resistance to herbivory in the field, we quantified physical damage from chewing insects on all individuals one week before herbicide application. Thus, herbivory estimates were determined for all plants but only in the absence of herbicide. To do this, we selected 3 leaves at random from each individual (620 individuals x 3 leaves = 1,860 total leaves examined) and photographed them completely attached to the plant in the field. We used the imaging software, LeafByte (Getman-Pickering et al 2020) to calculate the area of leaf tissue consumed by herbivores. We edited the photographs by cropping and removing background noise, then uploaded them into LeafByte. To obtain the total leaf surface area along with total chewing herbivory damage, we standardized an 8cm x 8cm area around each image allowing the software to accurately capture measurements.

### Operational Definition of Herbicide and Herbivory Resistance

We define herbicide resistance operationally as 1 minus the proportion of leaves exhibiting damage from the herbicide. Similarly, we define herbivory resistance as 1 minus the average amount of chewing damage across each of the 3 leaves per individual.

## Data Analysis

### Growth room study

#### Assay for herbicide resistance

We conducted all statistical analyses in R (version 3.4.2, R Development Core Team). A dose-response curve was fitted to the survival data for each of the populations, using the method of generalized log-logistic models with the binomial data implemented by the drc package (Ritz et al 2015). We estimated the logarithm of the dose which killed 50% of the plants for each population using the dose.p function from the MASS package (Venables and Ripley 2002). To uncover if there was genetic variation for herbicide resistance (measured as biomass) by treatment in the growth room, we fit the following mixed linear model using lmer function of the lme4 package (Bates et al. 2015):

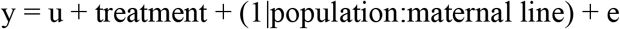

where herbicide resistance was the response variable, u is the intercept or mean of the model, treatment was a fixed effect, maternal line nested within population was a random effect, and e is the error term. We first standardized biomass within herbicide treatments by the control and the significance of effects was determined using the lmerTest package (Kuznetsova et al 2017).

Resistance was log + 1 transformed to meet the assumptions of the model.

### Field experiment

#### Correlations and variation among trichome traits

To determine if trichome traits were correlated, we performed Pearson’s correlation tests on the raw values of each of our trichome traits (single, branched, peltate, capitate, and total density).

To test for genetic variation for trichome traits, we fit the following model for each trait:

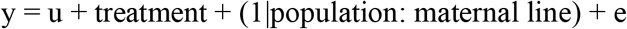

where each respective trichome trait was the response variable, u is the intercept or mean of the model, treatment was a fixed effect, maternal line nested within population was a random effect, and e is the error term. For each trichome trait, we removed the effects of block by performing a regression of the trait on block, and then used the residuals in each analysis of variance. We determined the significance of predictor variables using the stats package (Chambers and Hastie 1992) to generate *F*-statistics for the fixed effects. We used the lmerTest package (Kuznetsova et al 2017) to perform a log-likelihood ratio test for each random effect.

#### Trichome relationship to herbicide and herbivory resistance

Because the proportion of branched trichomes and capitate trichomes exhibited significant genetic variation (see Results) and because total trichome density is the most conventional trichome trait tested in the literature, we elected to perform the remaining analysis on these three traits. To determine if trichomes influence herbicide resistance in the field, we performed analysis of variance (ANOVA) by fitting a model for herbicide resistance:

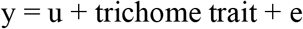

where, y, the response variable, is herbicide resistance, trichome trait (proportion of branched, capitate, or total density) as the dependent variable, and e is the error term. We determined the significance of the predictor variables using *F*-statistics for trichome characteristics using the stats package described above.

We performed a similar analysis of variance (ANOVA) to determine whether patterns of herbivory resistance were altered by trichome phenotype. We fit regressions of herbivory resistance on the trait in question:

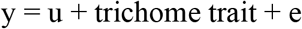

Prior to analysis, the herbivory was log transformed to improve normality. Because herbicide damage was only captured in the herbicide treatment and herbivory estimates were collected prior to herbicide application, treatment was excluded from the models. Preliminary analyses also indicated that there were not significant maternal line effects. We thus elected to exclude these effects from our final models.

#### Phenotypic Selection

We performed phenotypic selection analysis (Lande and Arnold 1983) to determine if herbicide alters selection on trichome traits. We estimated differentials to measure total selection on the proportion of branched trichomes and trichome density and gradients to measure direct selection on each. Relative fitness was calculated as the final seed count per individual divided by mean seed count for each treatment. Phenotypic traits were standardized to a mean of zero and variance of one prior to analysis. We estimated selection differentials (*S*) by performing univariate regressions of relative fitness on each trait separately as a measurement of both indirect and direct selection acting on each trait. We estimated linear selection gradients (*β*) using models only containing the linear terms as a measurement of direct selection on each trichome trait. Non-linear selection gradients (*γ*) were estimated using a full model containing linear terms, quadratic terms, and the cross-products terms of the focal trichome traits. By doubling the quadratic regression coefficients, we were able to examine the potential of quadratic (disruptive or stabilizing) selection or correlative selection.

Moreover, we used an ANCOVA to determine whether herbicide application significantly altered the pattern of selection on differentials. To do this, we performed a univariate regression of relative fitness on the trait in question, treatment, and their interaction, in which a significant interaction would indicate that herbicide significantly alters selection. We similarly tested whether linear and quadratic selection were significantly altered by treatment.

## Results

### Growth room study

*Assay for herbicide resistance–*Analysis of dose response curves revealed that the median lethal dosage (LD50) of glyphosate among populations was 740.63 g/ha ai, which is below the recommended field dose of 1,000 g/ha ai. We detected evidence of population differences in standardized biomass post-herbicide application (χ^2^ = 5.01, p = 0.025, SFigure 2B), indicating significant variation for glyphosate susceptibility. We found that the highest LD50 was population 4 with 1000 g/ha ae, which notably is the recommended field dose of this species, indicating reduced susceptibility may be evolving in this species (SFig 1A). Notably, however, we did not find evidence of maternal line variation for glyphosate resistance (χ^2^ = 0.00, p = 1.000).

### Field experiment

#### Correlations and genetic variation among trichome traits

We found that total trichome density was positively correlated with all trichome types expect for capitate trichomes (branched: r = 0.72, p < 0.001; single: r = 0.71, p < 0.001; peltate: r = 0.55, p < 0.001). We also found that peltate trichomes were positively correlated with single trichomes (r = 0.42, p < 0.001) and branched trichomes (r = 0.26, p = 0.002, SFigure 3).

We examined the potential for genetic variation in trichome traits to determine which traits may evolve in response to selection from the herbicide. We detected significant maternal line variation for proportion branched (χ^2^ = 3.96, p = 0.047, STable 1) and capitate trichomes (χ^2^ = 3.91, p = 0.048, STable 1) but not the other trichome types/traits (single: χ^2^ = 2.82, p = 0.092; peltate: χ^2^ = 0.75, p = 0.387; density: χ^2^ = 1.02, p = 0.313; STable 1).

#### Trichome relationship to herbicide and herbivory resistance

We examined the potential for a relationship between trichome traits and the level of herbicide resistance to determine if either the density of trichomes, the proportion of branched or capitate trichomes on the leaf surface may act as an herbicide resistance trait. We found that the proportion of branched trichomes was positively correlated to the level of herbicide resistance (F = 5.87, p = 0.019; Fig. 2), but that total trichome density (F = 0.00, p = 0.956; Fig. 2) and the proportion of capitate trichomes (F = 0.13, p = 0.719; Fig. 2) showed no such relationship. We found that the proportion of branched trichomes had a positive effect on herbivory resistance (F = 10.20, p = 0.002, Fig. 2). Capitate trichomes (F = 0.03, p = 0.235, Fig. 2) and total density of trichomes (F = 0.22, p = 0.789, Fig. 2) were not correlated to our measure of herbivory resistance.

**Figure 2.**
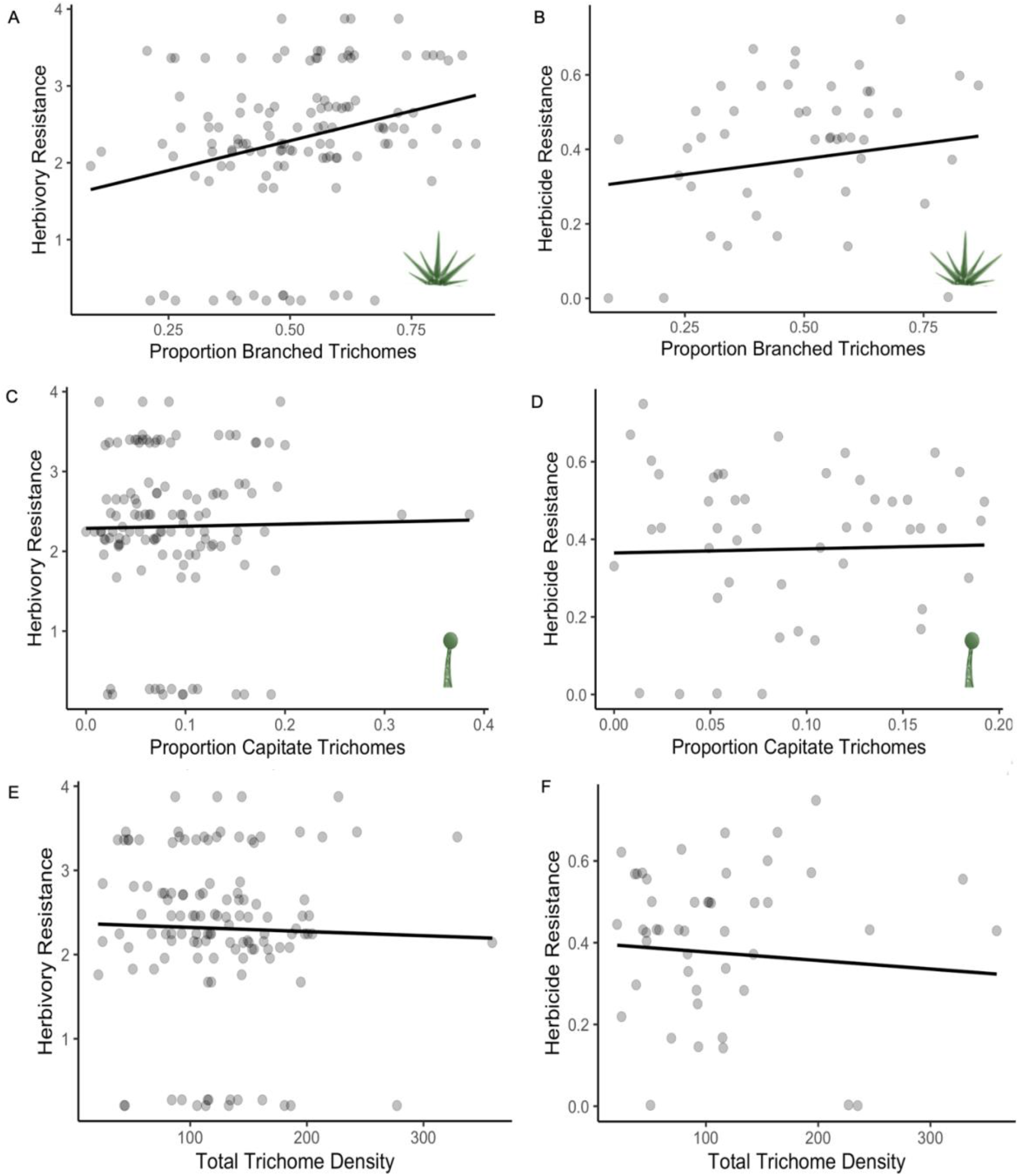
Relationships between the proportion of each trichome type or trichome density with resistance to herbivory and herbicide. A) The relationship between the proportion of branched trichomes and herbivory resistance (F = 10.20, p = 0.002) and B) proportion branched trichomes and herbicide resistance (F = 5.82, p = 0.019). C) The relationship between the proportion of capitate trichomes and herbivory resistance (F = 0.03, p > 0.05) and D) proportion of capitate trichomes and herbicide resistance (F = 0.64, p > 0.05). E) The relationship between total trichome density and herbivory resistance (F = 0.22, p > 0.05) and F) total trichome density and herbicide resistance (F = 0.09, p > 0.05). Datapoints represent individual plants; herbivory resistance was measured as log-transformed 1 – amount of chewing herbivory damage, and herbicide resistance was measured as 1 – proportion of leaf yellowing.

#### Phenotypic selection

We examined phenotypic selection differentials in both the control and herbicide treatments to determine if the pattern of selection on trichome traits was altered by the presence of herbicide. For the proportion of branched trichomes, the ANCOVA revealed that herbicide application significantly altered the pattern of selection (F = 8.20, p = 0.005, Table 1, Fig 3), with evidence for positive selection on the proportion of branched trichomes in the presence of herbicide (*S* = 2.05, p = 0.045, Table 1, Fig 3) and evidence for negative selection on this trait in control conditions (*S* = −1.98, p = 0.052, Table 1, Fig 3). Under control conditions, trichome density was under positive selection (*S* = 2.62, p = 0.011, Table 1, Fig 3), whereas there was no evidence of selection in the presence of herbicide (*S* = 1.77, p = 0.082, Table 1, Fig 3). There was no evidence of selection acting on the proportion of capitate trichomes in the presence (*S* = −1.28, p = 0.207) or absence of herbicide (*S* = −1.60, p = 0.112).

**Table 1.** Total selection (*S*) on velvetleaf trichome traits (proportion branched, proportion capitate, and total trichome density). Treatments are the absence (control) and presence of herbicide. Shown are selection differential values, standard errors, and p-values for traits in each treatment. The F-value indicates the treatment by trait interaction from an ANCOVA, and significant effects are indicated in bold.

| Phenotypic Selection (*S*) Differentials |  |  |  |  |  |  |  |
| --- | --- | --- | --- | --- | --- | --- | --- |
| Trait | Control |  |  | Herbicide |  |  | F-values (from ANCOVA) |
|  | *S* | SE | p-value | *S* | SE | p-value |  |
| Proportion Branched | **-1.98** | **0.10** | **0.052** | **2.05** | **0.22** | **0.045** | **8.20** |
| Proportion Capitate | -1.60 | 0.09 | 0.11 | -1.28 | 0.29 | 0.207 | 0.77 |
| Total Density | **2.62** | **0.13** | **0.011** | 1.77 | 0.20 | 0.082 | 0.01 |

**Figure 3.**
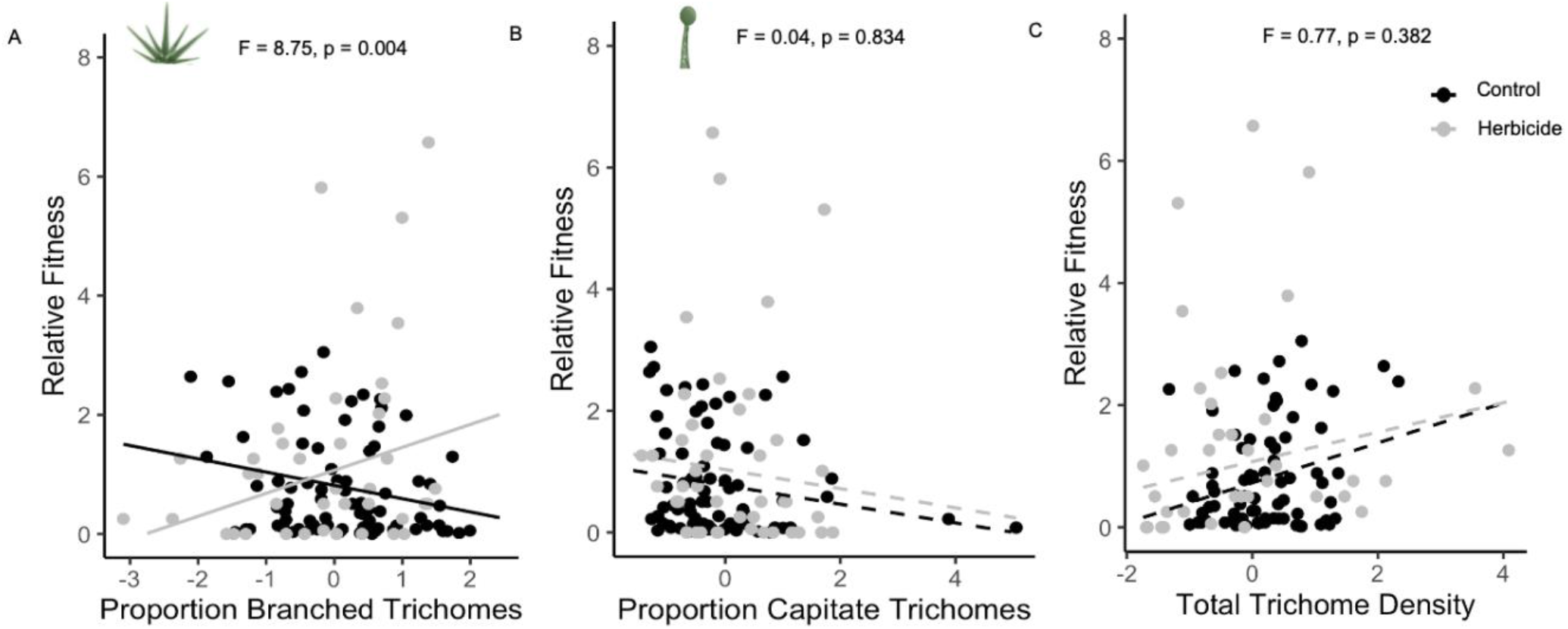
The relationship between relative fitness and A) the proportion branched trichomes, B) proportion capitate trichomes, C) and total trichome density in the presence (grey) and absence (black) of herbicide in *Abutilon theophrasti* in field conditions. Solid lines represent significant selection differentials in each treatment environment and F-statistics show the treatment by trait interaction from the ANCOVA.

Linear selection gradients were similar to the selection differentials. We identified linear selection favoring increased trichome density in the absence of herbicide (β = 2.06, p = 0.043, Table 2) but not in the presence of herbicide (β = 1.38, p = 0.06, Table 2). We also found that the proportion of branched trichomes was under positive selection in the presence of herbicide (β = 2.16, p = 0.036, Table 2), and that herbicide exposure significantly altered the pattern of selection on the proportion of branched trichomes (β = −1.96, p = 0.053, ANCOVA: F = 8.75, p = 0.004, Table 2). We found no evidence of selection on the proportion capitate trichomes in either the presence (β = −0.34, p = 0.622, Table 2) or absence of herbicide (β = −0.86, p = 0.393, Table 2). Overall, these results indicate that trichome density and the proportion of branched trichomes are under direct selection and not simply linked to other traits selected upon. We did not identify evidence of quadratic selection acting upon branched trichomes, trichome density, or their interaction in the presence or absence of herbicide (STable 2).

**Table 2.**
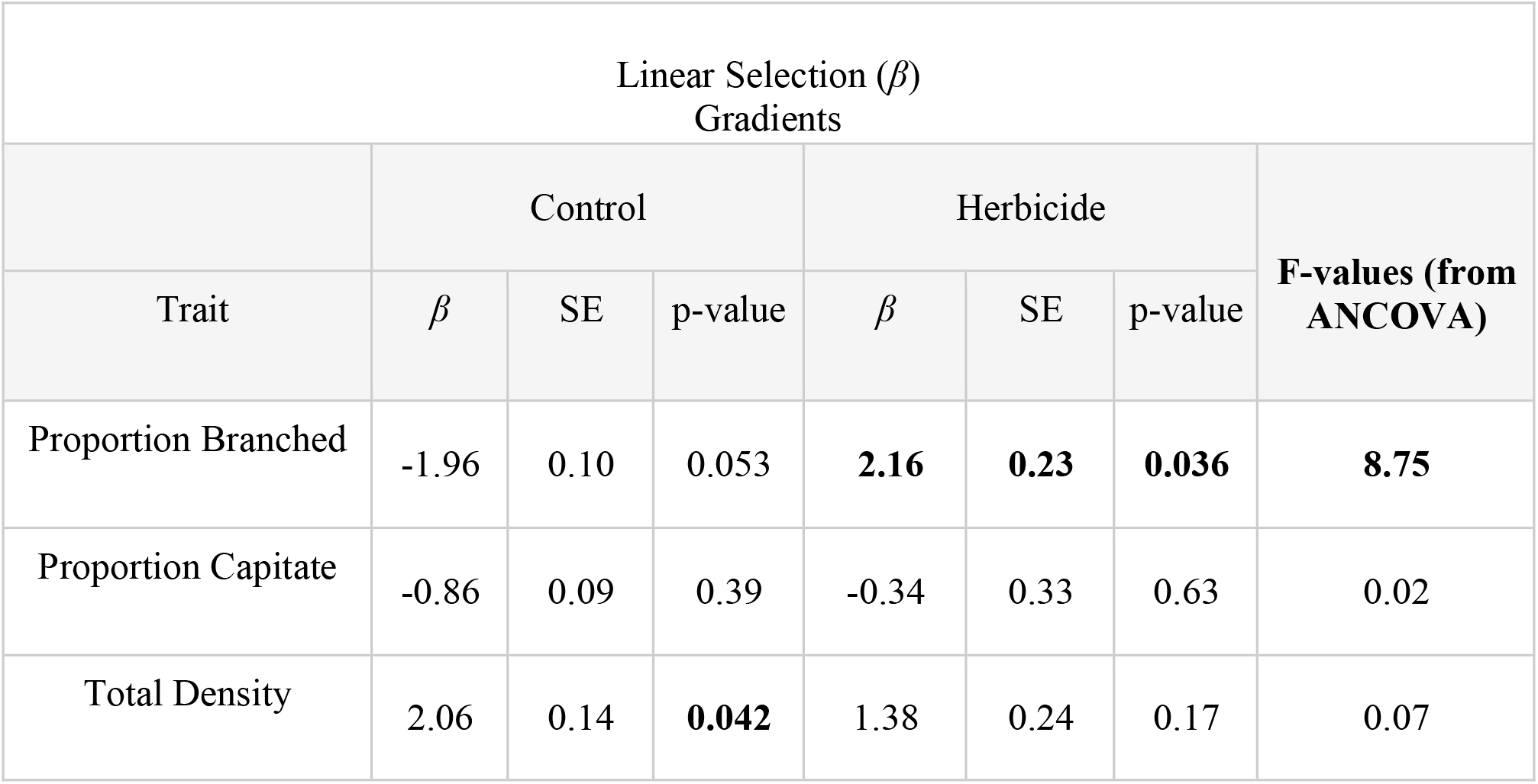
Direct (multivariate) selection acting on trichome traits (proportion branched, proportion capitate, and total trichome density) in the absence and presence of herbicide. Shown are linear (*β*) gradient values, standard errors, and p-values in each treatment. F-values are from the ANCOVA analysis testing the effect of herbicide treatment on selection gradients. Significant effects are indicated in boldface.

## Discussion

The goal of our study was to determine if trichome traits exhibit dual or conflicting roles in plant defense when faced with different forms of damage using the common weed *Abutilon theophrasti*. We found positive correlations between the proportion of branched trichomes and both herbicide and herbivory resistance, indicating that trichomes can serve a dual defensive role by acting as both an herbivory and herbicide resistance trait. We also showed that herbicide acts as an agent of selection on the proportion of branched trichomes. Our findings expand current knowledge of how plant trichome phenotypes may evolve in response to multiple stressors and highlight the importance of studying trichome traits beyond that of trichome density.

### Branched trichomes in plant resistance

Though the relationship between trichomes and herbivory resistance has been well studied, to our knowledge, this is the first time that trichomes have been considered as a potential herbicide resistance trait. While we do not know how trichomes might function as an herbicide resistance trait, we hypothesize they may either store the herbicide or reduce its contact with the leaf surface. For example, it is possible that herbicide might be compartmentalized into trichome cell walls and vacuoles, like the compartmentalization of heavy metals into the trichomes of heavy-metal tolerant serpentine plants (Cobbett 2000, Küpper et al 2000, Broadhurst et al 2004,

Marmiroli et al 2004). In support of this idea, branched trichomes have a greater vacuole volume than other trichome types (Calvert et al 1985) as the vacuole occupies 90% - 95% of the total trichome volume (Gutierrez-Alcalcaet at al 2000). Alternatively, increased amounts of branched trichomes on leaves may provide a more complex network of interweaving appendages, acting as a cohesive umbrella of sorts that holds herbicide droplets on the upper epidermal layer due to surface tension and reducing absorption into the plant altogether (*i*.*e*., providing a barrier that reduces herbicide damage in the plant body, Baucom 2019).

Our study also demonstrates that the proportion of branched trichomes is correlated to chewing herbivory damage such that a higher proportion of branched trichomes is positively related to higher herbivory resistance. These results align with work in cotton showing that accessions with the highest amount of branched trichomes have the greatest resistance to spider mites (Kamel and Elkassaby 1965). Somewhat surprisingly, we did not find an association between chewing herbivory resistance and total trichome density in velvetleaf; in this way our results differ from findings from other plant species, in which individuals with a greater density of trichomes evade herbivory damage more successfully than individuals with lower densities of trichomes (Doss et al 1987, Webster et al 1994, Romeis et al 1999).

There are several possible ways in which a higher proportion of branched trichomes may mitigate damage from a range of herbivorous species. Trichomes make it difficult for the insect to access the plant body, may cause entrapment and inflict injury (Dalin et al 2008, Sxyndler 2013), and inhibit herbivore growth and development via post-ingestive effects (Shanower 2008, Kariyat et al 2017). Branched trichomes in particular cover a larger surface area than other trichome shapes and may either entrap or impede insects. In our field study, Japanese emerald beetles and grasshoppers were the most common herbivorous insects feeding on velvetleaf, thus, these large insect herbivores may prefer plants with a lower proportion of branched structures to avoid their negative effects.

### Dual defensive function

Environmental heterogeneity is thought to be a primary mechanism of maintaining phenotypic variation (Hedrick 1986). Under this premise, we might expect defense traits that contribute to resistance against traditional forms of damages such as herbivory in one environment to differ or conflict with traits that contribute to resistance against novel forms of damage in another environment such as herbicide exposure. Instead, we find that branched trichomes are serving a dual defensive role in that they are mutually contributing to herbivory resistance and herbicide resistance. An important caveat to our study is that we did not reduce or remove herbivory when examining the potential for a relationship between trichome traits and the level of herbicide resistance. Thus, if herbivory induces an increase in the proportion of branched trichomes prior to herbicide exposure, the positive relationship we identified between the level of resistance and the proportion of branched trichomes may simply be due to induced effects from herbivore damage. While this is an important caveat of our study, a trait may offer a fitness advantage and be classified as defensive without identifying its primary function (Strauss and Agrawal 1999). In *Verbascum thapsus*, for example, trichomes on younger leaves have been identified as a defense against grasshopper herbivory but also protects against water loss in high temperatures (Woodman and Fernandes 1991). In the context of agricultural fields, plants would experience herbivory and herbicide spraying, such that even if herbivory induces a higher proportion of branched trichomes, the higher proportion present on leaves as a result would still be functionally relevant for subsequent herbicide exposures.

### Adaptive significance of trichome traits

Given the importance of trichomes in mitigating effects from numerous abiotic and biotic stressors, it is important to understand the processes that promote variation in trichome traits, since selection imposed by one stressor may impact adaptation to another. We examined the potential for selection on trichome traits and focused on the proportion each of branched and capitate trichomes, since these traits exhibited genetic variation. Although we did not detect significant variation in trichome density for the populations in this study, we included analysis of trichome density since density is known to be a highly variable character in several other species (Heinz and Zalom 1995, Roy et al 1999, Valverde et al 2001).

Interestingly, our analyses revealed that selection on trichome traits differed by herbicide treatment. In the absence of herbicide, we found evidence of negative selection on branched trichomes and in the presence of herbicide we uncovered positive selection on branched trichomes. This result indicates that herbicide acts as an agent of selection on the proportion of branched trichomes. Alternatively, in the absence of herbicide, we found positive selection on trichome density but no evidence of selection on this trait in the presence of herbicide, suggesting that in non-herbicide environments, trichome density may increase when plants experience herbivory. We note, however, that we did not identify genetic variation in trichome density in our study population, meaning that we would not expect a response to this selection. Further, for both trichome traits, we found that the selection gradients showed the same results as the differentials indicating that selection is directly acting on these traits in the presence of herbicide, and there were no indirect effects acting on either trait. Finally, there was no evidence of correlative selection acting upon trichome density and the proportion of branched trichomes in either environment.

## Conclusion

A major objective of plant evolutionary ecology has been to understand how defense strategies differ and are maintained over time and space (Feeney 1976). Though it has been proposed that the evolution of specific assortments of traits may be associated with particular environmental factors (Grime 1977, Chapin et al 1993), this framework has rarely been applied to explore the potential adaptive significance of trichome composition in complex environments (Escobar-Bravo et al 2019, Karabourniotis et al 2020). We explored how the proportion of trichome types, beyond the traditional classification of glandular and non-glandular, can provide novel insight into plant defense strategies when faced with conventional and modern damaging agents in agroecosystems. Moreover, in identifying variation of trichome landscapes among individuals and populations, we present an approach that may help us to better understand the evolution of plant defense.

Overall, our study reveals that branched trichomes contribute both to herbicide resistance and herbivory resistance in *A. theophrasti*. The results also suggest plants exposed to glyphosate could become better defended against chewing herbivores, since glyphosate imposes positive selection on the proportion of branched trichomes a plant exhibits, which in turn increases resistance to herbivory. If such evolution occurred, it could influence herbivore population growth, habitat choice, and subsequent multi-trophic interactions. As such, future studies should investigate whether trichome composition correlates with distance from agricultural fields and herbivore abundance over long timescales. In summary, our findings are that changes in trichome composition against one stressor may impact defense against other stressors. These changes could also have cascading effects upon other aspects of the food web. Understanding the functionality of trichomes as structural defensive traits may help us better predict ecological evolution of communities most responsive to global environmental change.

## Acknowledgments

Many thanks to Kaira Liggett, Shantrell Tremmell, and Alanna Miyashiro who helped with field maintenance and with capturing pictures for herbivory estimates. Raj Gautam and Jordan Herman who helped collect and count seeds. Michael Palmer and Jeremy Moghtader at Matthaei Botanical Gardens for helping to till the field plot and apply the herbicide. Gregg Sobocinski at the University of Michigan Imaging Core for assistance with capturing trichome images. The manuscript was improved by insightful comments from Ines Ibanez, Deborah Goldberg, D. André Green, Meghan Duffy, Marjorie Weber, Bruce Martin, Sylvie Martin-Eberhardt, Mia Howard, Abrianna Soule, Rosemary Glos, and Amana Vieria da Silva. This research was supported by the Ecology and Evolutionary Biology Department at the University of Michigan.

## Notes

### Competing Interest Statement

The authors have declared no competing interest.

