## Supplemental Figure 1 for "The double life of trichomes: understanding their dual role in herbivory and herbicide resistance"

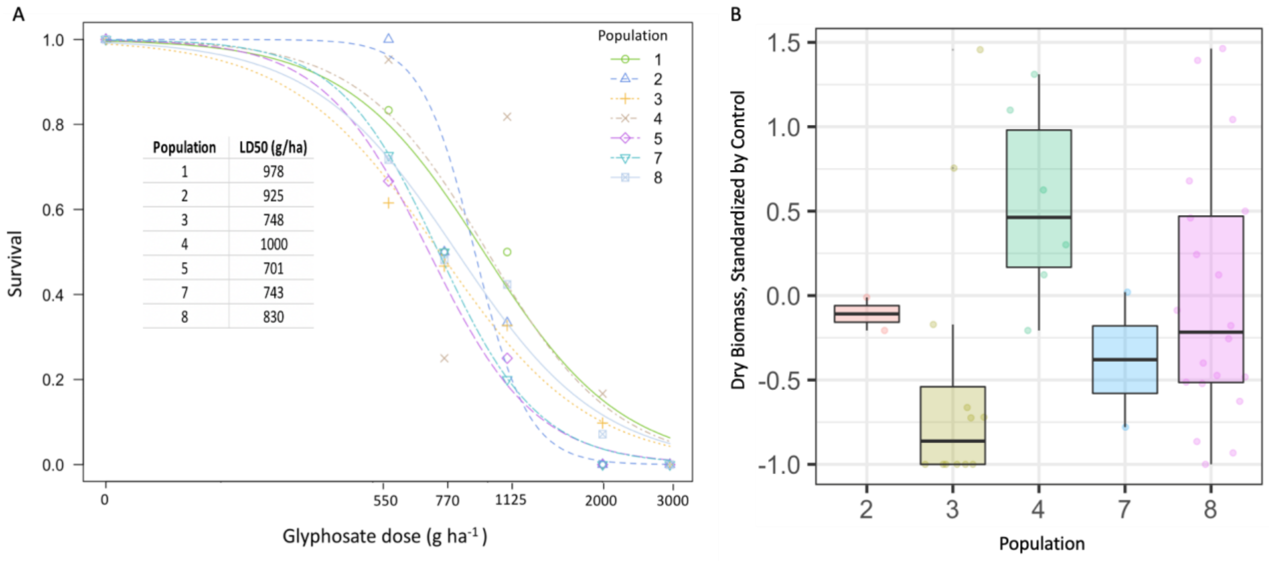


SFigure 2A. Dose-survival response relationships of 8 *Abutilon theophrasti* MI populations after treatment with glyphosate at 550g/ha ai, 770g/ha ai, 1125g/ha ai, 2000 g/ha ai, and 3000 g/ha ai measured in the growth room experiment. Shown are median lethal dosage (LD50) among the populations sampled. B. Bar graph shows the variation of dry biomass at the dose 550g/ha ai standardized by the controls across populations 2, 3, 4, 7, and 8. We determined the fixed treatment effects ($F = 51.92$, $p > 0.001$) using Type III Sums of Squares and the significance of the random population effects using chi-squared test ($\chi^2 = 5.01$, $p = 0.025$).
