## Supplemental Figure 2 for "The double life of trichomes: understanding their dual role in herbivory and herbicide resistance"

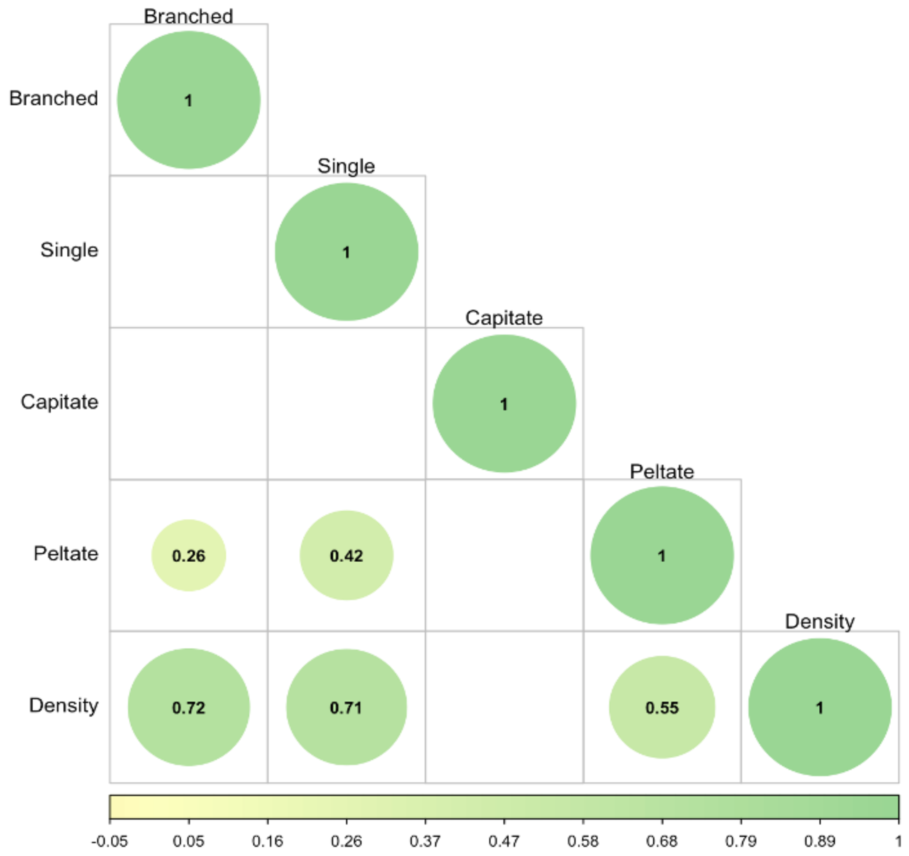


SFigure 3. Pearson correlations describing the correlation between trichome traits represented by color gradients of green to yellow (1 to -1). The presence of a circle indicates a significant trait correlation. Circle size indicates p-value.
