## Supplemental Table 1 for "The double life of trichomes: understanding their dual role in herbivory and herbicide resistance"

| | Maternal Line | | | Population | | |
|---|---|---|---|---|---|---|
| | **$\chi 2$** | df | p-value | **$\chi 2$** | df | p-value |
| Branched | **3.96** | **1** | **0.047** | 0.00 | 1 | 1.000 |
| Single | 2.82 | 1 | 0.092 | 0.00 | 1 | 1.000 |
| Capitate | **3.19** | **1** | **0.048** | 0.00 | 1 | 1.000 |
| Peltate | 0.75 | 1 | 0.387 | 0.00 | 1 | 1.000 |
| Density | 1.02 | 1 | 0.313 | 0.39 | 1 | 0.531 |

STable 1. Results from a test for genetic variation using chi statistics ($\chi^2$) values showing the effects of maternal line and population variation on trichome traits (proportion branched, single, capitate, peltate, and trichome density) captured in the field. Significant effects are indicated in boldface.
