## Supplemental Table 2 for "The double life of trichomes: understanding their dual role in herbivory and herbicide resistance"

| Quadratic Selection ($\gamma$) Gradients | | | | | | | |
|---|---|---|---|---|---|---|---|
| | Control | | | Herbicide | | | **F-values (from ANCOVA)** |
| Trait | $\gamma$ | SE | p-value | $\gamma$ | SE | p-value | |
| Proportion Branched | 1.89 | 0.54 | 0.349 | -0.26 | 1.94 | 0.898 | 0.01 |
| Proportion Capitate | -0.97 | 0.28 | 0.63 | 0.59 | 2.56 | 0.769 | 0.02 |
| Total Density | 0.38 | 0.92 | 0.85 | 0.27 | 1.75 | 0.895 | 1.24 |
| Proportion Branched x Proportion Capitate | -1.34 | 0.17 | 0.51 | -1.67 | 3.12 | 0.41 | 0.13 |
| Proportion Capitate x Density | -1.22 | 1.33 | 0.55 | 2.32 | 4.01 | 0.257 | 0.52 |
| Proportion Branched x Density | 0.68 | 0.98 | 0.737 | 0.62 | 3.81 | 0.76 | 0.15 |

STable 2. Direct (multivariate) selection acting on trichome traits (proportion branched, proportion capitate, density, and their interactions) in the absence and presence of herbicide. Shown are the quadratic ($\gamma$) gradient values, standard errors, and p-values in each treatment. F-values are from the ANCOVA analysis testing the effect of herbicide treatment on selection gradients. Significant effects are indicated in bold.
